# Diversity of arbuscular mycorrhizal fungi under the rhizosphere soil of different cropping systems surrounding Hawassa town, South Ethiopia

**DOI:** 10.1101/2023.11.01.565191

**Authors:** Girma Zeleke, Beyene Dobo, Fassil Asefa

**Affiliations:** College of natural and computational science; Department of Biology; Hawassa Univeristy, Ethiopia; Department of microbial, cellular and molecular biology, Addis Ababa University, Addis Ababa, Ethiopia

**Keywords:** *Arbuscular Mycorrhizal Fungi*, *Colonization*, *Diversity*, *Hawassa*, *Spore density*

## Abstract

This research was conducted to study the diversity of Arbuscular Mycorrhizal Fungi (AMF) in the rhizosphere of different plants at the vicinity of Hawassa city, Southern Ethiopia. The rate of root colonisation and diversity of AMF in the rhizosphere soils of permanent crops, major annual crops, forest areas, and open grazing fields were investigated. It discovered 928 spores of 23 distinct AMF morpho-species in 12 AMF genera and 12 annual and perennial crops. The AMF genera identified were: Acaulospora, Cetraspora, Claroideoglomus, Dentiscutata, Diversispora, Funneliformis, Gigaspora, Glomus, Racocetra, Rhizophagus, Sclerocystis, and Scutellospora. In tomatoes grown inorganically, a compromised species richness may result from the extensive use of agrochemicals. Further research into the effects of agricultural inputs on the subsurface microbial population may be necessary, as this is outside the purview of this publication. When compared to all other land uses, the AMF beneath the rhizosphere soil of Eucalyptus trees has the largest biomass, with spore density of 1907.4±0.404 spores 100g-1 of dry soil. The lowest AMF biomass has been recorded in the rhizosphere soil of Mango tree, with spore density of 260.1±0.121 spores 100g^-1^ dry soil. The results of this study show that the mycorrhizal colonisation and spore density of the plants under investigation are decreased by monocropping, intense agricultural practises, and the application of inorganic fertilisers. Thus, it is thought that conservation agriculture, together with the development of plant species consortiums in agricultural polts, will preserve the biological diversity of Arbuscular mycorrhizal fungi and the integrity of the environment.

**Significance of the study to the field:**

- plant species diversity is dependent on arbuscular mycorrhizal fungi diversity and density
- AMF Favours environmental resilience, carbon sequestration, and improves soil structure
- Application of inorganic agricultural inputs decrease, AMF diversity
- AMF diversity is affected by land use change, environmental pollution and climate change

## 1. INTRODUCTION

Arbuscular Mycorrhizal Fungi (AMF) is a group of obligate bio-trophs, to the extent that they must develop a close symbiotic association with the roots of a living host plant in order to grow and complete their life cycle [1]. The term “mycorrhiza” literally derives from the Greek ‘mykes’ and ‘rhiza’, meaning fungus and root, respectively. AMF can symbiotically interact with more than 80% of the plants on the Earth. Molecular DNA sequencing-based analyses have recently contributed to a great extent by shedding light on a previously unseen and profound diversity within this phylum [2].

AMF are found in the roots of about 80-90% of plant species (mainly grasses, agricultural crops and herbs) and exchange benefits with their partners, as is typical of all mutual symbiotic relationships [3]. They represent an interface between plants and soil, growing their mycelia both inside and outside the plant roots. AMF provide the plant with water, soil mineral nutrients (mainly phosphorus and nitrogen) and protection from pathogens. In exchange, photosynthetic compounds are transferred to the fungus [4]. AMF are probably the most ubiquitous fungi in agricultural soils, accounting for 5-36 % of the total biomass in the soil and 9-55 % of the biomass of soil microorganisms [5].

The populations of AM fungi is greatest in plant communities with high diversity such as tropical rainforests and temperate grasslands where they have many potential host plants and can take advantage of their ability to colonize a broad host range (Smith and Read, 2002). There is a lower incidence of mycorrhizal colonization in very arid or nutrient-rich soils. Mycorrhizas have been observed in aquatic habitats; however, waterlogged soils have been shown to decrease colonization in some species [6].

However, the effects of factors such as plant community in the AMF community composition are less clear [7]. In general, infectivity and diversity of AMF communities is often reduced in disturbed habitats such as agro-ecosystems or post-mining sites [8]. The specificity, host range, and degree of colonization of mycorrhizal fungi are difficult to analyze in the field due to the complexity of interactions between the fungi within a root and within the system. There is no clear evidence to suggest that arbuscular mycorrhizal fungi exhibit specificity for colonization of potential AM host plant species as do fungal pathogens for their host plants [6].

Though they are few, scholars in Ethiopia indicated that the difference in the cropping system affected the diversity of AMF. For instance, according [9], land use types drastically affected AMF colonization and AMF diversity in a dryland agroforestry system in the central part of Ethiopia. Besides, the study carried out by [10, 11] revealed that the number of spore count was significantly higher under the canopy of trees than outside the canopy. Meanwhile the unpublished study conducted in Northern part of Ethiopia, Jabi Tehnan woreda western Gojam *by* [12] reported that spore density of the different cropping systems varied significantly within and between land use types. Moreover, study findings of [13] confirm distinct fungal communities associated with the diverse tree species.

Studies done by [14], in Sidama region, South Ethiopia, revealed the role of the physicochemical nature of the soil in AMF abundance and density. In that the soil with pH of 6.18-6.28 and texture of clay loam and sandy loam favored mycorrhizal development. Besides, [14] reported favorable AMF growth at moderate to low concentration of P and N.

Information regarding the diversity of AMF under the rhizosphere soil of various cropping system in Ethiopia is very minimal. Particularly, very little or nothing is known regarding the AMF diversity associated with the cropping system and soil in Hawassa area. Nonetheless, trends of AMF diversity particularly associated with the use of inorganic fertilizers are also seeking attention in this specific area.

This kind of study would attribute to food security and agricultural development agendas of the nation. By 2050, global agriculture will have the task of doubling food production in order to feed the world [15]. At the same time, dependence on inorganic fertilizers and pesticides must be reduced. For these reasons, significant advances in AMF research are needed to allow their stable use in agriculture. Their application and synergistic combination with other functionally efficient microbial consortia that include PGPR (Plant Growth Promoting Rhizobacteria), saprophytic fungi and other helper microorganisms will help farmers develop a more sustainable cropping system [16]. Hence, findings of the study substantiate knowledge regarding the diversity and abundance of AMF across the different cropping systems around Hawassa city, Sidama regional state of Ethiopia.

## 2 Materials and Methods

### 2.1 Study area Description

The study has been undertaken at randomly selected agricultural plots and open grass land allotted for livestock grazing. All the farm plots and the grass land located at the out skirt of Hawassa area. Hawassa is located in the north-eastern part of the SNNP region and bounded by Oromiya region in the North, Shebedeno district in the south Wondogenet district in the east and Lake Hawassa in the west. Its geographic location lies between 6^0^91′ and 7^0^10′ North latitude and 38^0^41′ and 38^0^55′ East longitude, with altitude ranges form 1501-2500masl. Total area of the Hawassa City is about 271 Km^2^. As per the 2014/15 statistical abstract of the SNNPR, the population of Hawassa city administration reached 355,610. Average population density of the city is 1,312 person per/ km^2^ of which 35% of the population represents the rural inhabitants, who mainly rely on agricultural activity to support their livelihood (SNNPR-BoFED, 2014/15).

The rural portion of the Hawassa administration covers about 11,940hectares. The land use of this rural area is characterized as: annul crops (2,409ha), permanent crops (3,360ha), grazing land (850ha), forest (1,156ha), potentially cultivable (275ha), uncultivable (165ha) and others (3,725ha). Mean annual rainfall of the area ranges from 801-1400mm. The mean annual temperature of the area ranges between 17-22.5°c [17].

### 2.2 Study design

Descriptive cross-sectional field survey is employed to understand the rate of root colonization and diversity of AMF under the rhizosphere soils of permanent crops, annual major crops, forest lands and open grazing fields.

### 2.3 Field Sample collection

Soil and root samples were collected to determine the AMF diversity and evaluate root colonization respectively. Sampling were carried out in randomly selected crop lands (organically cultivated tomato, inorganically cultivated tomato, Enset, Coffee, Maize, Eucalyptus, Wanza, Banana, Shewshewe (Oak), Mango, Pepper and an open grazing field. Sample collection was made from Feb-April 2018.

#### 2.3.1 Soil and Root Sample Collection

A composite soil of 300g, with in a depth of 4-20cm, was collected from the rhizosphere of each study plants (Table 1). The soil samples air-dried and packed individually in sterile polythene bags and stored at 4° C until processing for extraction and enumeration of AM fungal spores. A total of (12×3) soil samples werecollected. The soils samples collected from each sites were divided into three portions to enable three times replication and transported to Microbiology laboratory at Hawassa teacher training center. Recoveryof very fine roots of AMF, from the rhizosphere of eachstudy plantwas realized through the collection of fine root samples of 0.5mg (12×3) and it has been cut into 1cm pieces, washed with tap water,Put in a test tube and preserved in 50% ethanol and stored at 4°C for further analysis [18].

**Table 1:**
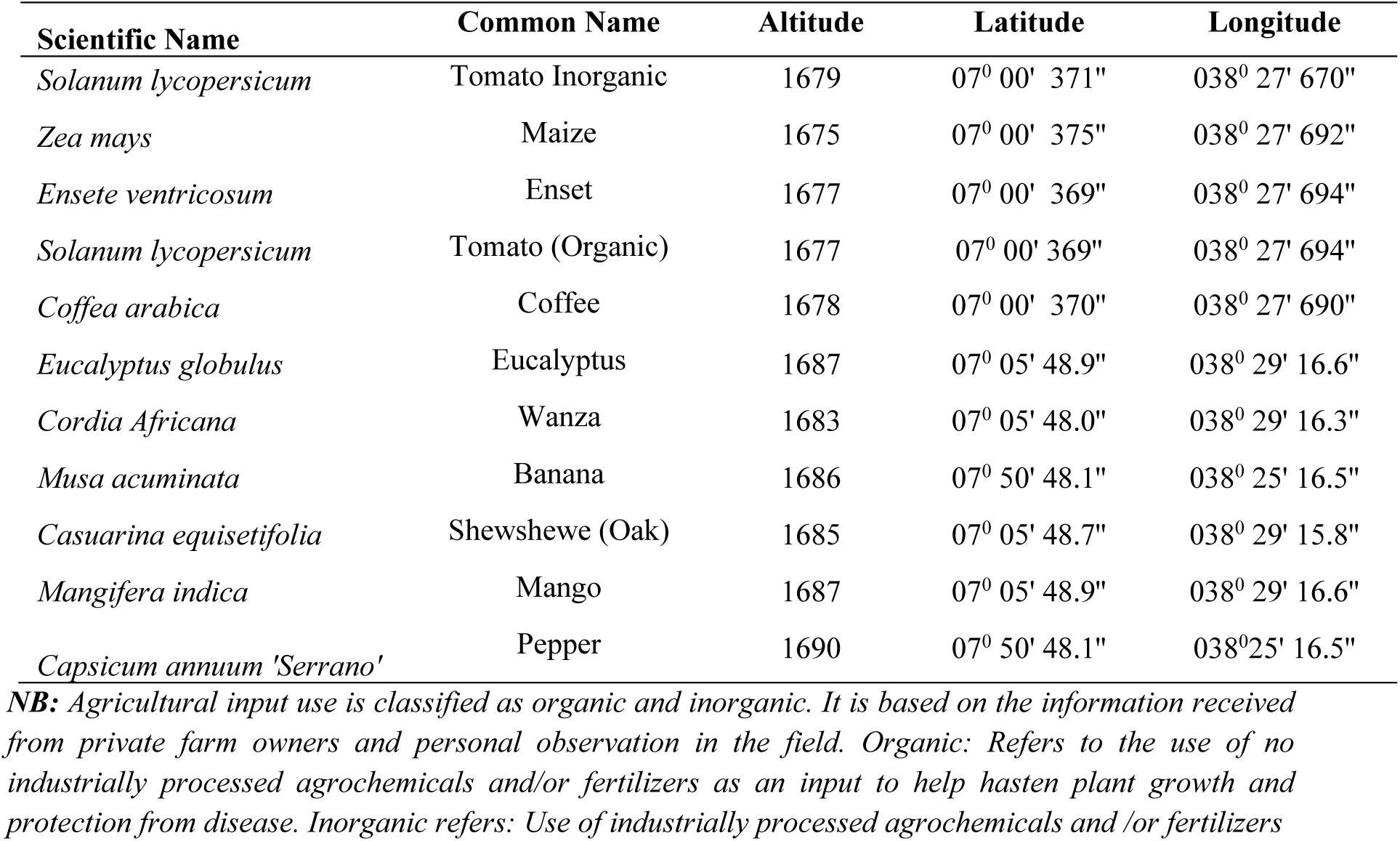
Georeferenced rhizosphere soil and root sampling points, Hawassa, Ethiopia.

### 2.4 Sample Analysis

Analysis of the rhizosphere soil and roots were made to investigate AMF diversity and estimate the rate of root colonization. It was an engagement fromMay-Aug, 2018.

#### 2.4.1 Investigation of AMF root colonization of study plants

AMF root colonization was assessed according to [19]. Root samples were washed several times with tap water and cleared in 10% (w/v) KOH by heating in a water bath at 90°C for 1-2 hrs and cooled at room temperature. After cooling, the root samples were washed 3-5 times with tap water, acidified in 1% HCl for 1hr and stained with 0.05% trypan blue and finally distained in acidic glycerol. The AM fungal structures were observed under a compound-light microscope (Olympus-bx 51) at 200 fold magnification. Fungal colonization were estimated using the magnified intersection method of [20] as total root length colonization RLC=100 [(G-N)/G], the percentage of root length colonized by arbuscules, arbuscular colonization AC = 100 (A/G) and the percentage of root length colonized by mycorrhizal vesicles, vesicular colonization VC = 100 (V/G). RLC, N, A, V and G are designated as RLC (total root length colonization), N (no fungal structure), A (arbuscules), V (vesicles) and G (total intersection) respectively. All were quantified by examining 100-150 intersections per sample.

#### 2.4.2 Identification and quantification of AMF spores within the rhizosphere soil of study plants

AMF spore diversity (species range and enumeration) work was determined rendering methodology by [21]. Accordingly, 100g of each soil sample were suspended into 2 liter container and mixed vigorously to free spores from the soil and roots. The supernatant was subsequently decanted through standard sieves (480, 106, 50 & 38μm) after having been intermittently centrifuged at 2000rpm for 5 minutes. The last pellet (38μm) were suspended in 60% sucrose solution and thoroughly mixed and centrifuged at 2000rpm for 1 minute to collect the spores. The spores and sporo-carps, from the upper solution of the centrifuged tube, were then be rinsed with tap water and transferred into plastic petri-dishes. Counting has been madeunder 4x stereomicroscope according to http://invam.caf.wvu.edu and spore density was expressed as the number of spores and sporo-carps per 100gof dry soil. Healthy looking spores was collected and mounted on slides with polyvinyl-lactic acid-glycerol (PVLG) to identify them into the representative morpho-species based on the descriptions of the International Culture Collection of Vesicular/Arbuscular Mycorrhizal Fungi (http://invam.caf.wvu.edu; 2006), and following descriptions by (Schenck and Perez,1990) using a compound light microscope (Olympus-bx 51) at X200 magnification.

Quantitative study or spore enumeration work was done according to INVAM, http://invam.caf.wvu.edu.as follow. First using a fine ruler, the diameter of the ocular field of the stereomicroscope at a magnification of 40X was determined (where spores can be easily distinguished from mineral particles and organic debris). The area of the spherical field was calculated at that magnification (40X). It was having a diameter of 5mm that the area of the field was 19.6mm^2^(radius 2.5mm). Plastic Petri dishes were used to count spores because the base of the plate is flat. Because the dish also is hydrophobic, enough water is added to have complete coverage of the base. Those dishes which were 85mm across were used. That is, the area of the base that was calculated to be 5672mm^2^. From this datum and the area of the ocular field (19.5mm ^2^), and total number of fields in the dish was: 5672/19.6. After that, the extracted spore suspension was added to a petri dish and then the dish was rotated randomly to spread out spores as evenly as possible. Finally, spores were counted in 40 fields randomly chosen over the area of the dish. Average number of spores per field was calculated and multiplied by 289 (#fields/dish).

#### 2.4.3 Determination of AMF diversity

The AMF communities from (12×3) sampling areas were detected and calculated based on the following parameters: Spore density (SD) wase xpressed as the number of AMF spores 100g^-1^soil. Species richness (S) was measured as the total number of morpho-species. The Shannon–Wiener index (H′) of diversity was calculated using the formula: Hꞌ= -Σ ((ni /n) ln (ni/n)) where: ni = number of individuals of species i and n = number of all individuals of all species. The Simpson’s dominance index (D) was calculated using the formula D = Σ (ni/n)2; Evenness (E) was calculated by dividing Shannon–Wiener diversity value by the logarithm of the species richness. These analyses were conducted using the software PAST3 (ver. 3.0). Isolation frequency (IF) was calculated as (the number of samples in which a given species was isolated/ the total number of samples) ×100%. Relative abundance of spores (RA) was calculated as (the number of spores in a given species / total number of spores) ×100%. The importance value (IV) was used to evaluate the dominance of AMF species based on IF and RA and was calculated as IV = (IF + RA)/2. An IV ≥ 50% indicates that a genus or species is dominant; 10% < IV < 50% applies to common genera or species; an IV ≤ 10% indicates that a genus or species is rare [22].

### 2.5 Data Analysis

Data analysis was carried out with SPSS software (version 21). Data on percentage of AM colonization was transformed by arcsine X^½^ and spore densities were transformed by log (x+1) to fulfil the assumption of normality and homogeneity of variances before analysis of variance [23]. Means given in tables were subject to one-way ANOVA to test the differences in AM colonization and spore density among the cropping plants of the fields. Mean separation was done by Duncan’s multiple-range test at the 0.05 level of probability.

## 3 Result and Discussion

### 3.3 Soil Physico-chemical analysis

The study on soil parameters show that the soil texture is of three categories; clay, sandy loam and sandy clay loam with optimum concentration of pH, EC, OC, Total nitrogen and available phosphorus (Table 2). The correlation analysis also shows positive association between spore density and percentage of OC and total Nitrogen was identified, yet the association was not statistically significant (r= 0.514, P<0.088 and r=0.513, P< 0.088 respectively). Similarly, spore density of AMF associate positively with that of OC and total Nitrogen (r=0.411 P<0.185 and r=0.409, P<0.187 respectively), nonetheless it was not statistically significant association.

**Table 2:** Physico chemical results of rhizosphere soil sampled from the study area, Hawassa Ethiopia.

| Crop | pH | Electric<br>Conductivity (ms) | Carbon<br>(%) | Total Nitrogen<br>(%) | Available P<br>(ppm) | Texture |  |  | Texture Class |
| --- | --- | --- | --- | --- | --- | --- | --- | --- | --- |
|  |  |  |  |  |  | Sand (%) | Clay (%) | Silt (%) |  |
| *Coffea arabica* | 7.58 | 553 | 1.56 | 0.134 | 10.4 | 59 | 22 | 19 | Sand Clay Loam |
| *Mangifera indica* | 7.34 | 565 | 1.17 | 0.101 | 11.6 | 55 | 24 | 21 | Sand Clay Loam |
| *Cordia africana* | 7.04 | 992 | 3.90 | 0.336 | 15.8 | 51 | 26 | 23 | Sand Clay Loam |
| *Zea mays* | 5.05 | 1413 | 1.95 | 0.168 | 11.2 | 69 | 20 | 11 | Sandy Loam |
| *Casuarina equisetifolia* | 6.41 | 332 | 1.17 | 0.101 | 10.2 | 37 | 46 | 17 | Clay |
| *Musa acuminata* | 5.11 | 1449 | 2.34 | 0.202 | 14.2 | 65 | 26 | 9 | Sand Clay Loam |
| *Solanum lycopersicum (org)* | 7.24 | 392 | 1.56 | 0.134 | 9.2 | 67 | 24 | 9 | Sand Clay Loam |
| *Eucalyptus globulus* | 6.26 | 715 | 1.56 | 0.134 | 9.0 | 69 | 24 | 7 | Sand Clay Loam |
| *Open Grass Land* | 7.32 | 210 | 0.78 | 0.067 | 8.8 | 53 | 32 | 15 | Sand Clay Loam |
| *Capsicum annuum* | 6.76 | 1843 | 1.56 | 0.134 | 9.6 | 57 | 28 | 15 | Sand Clay Loam |
| *Solanum lycopersicum (inorganic)* | 6.55 | 602 | 1.36 | 0.118 | 8.2 | 59 | 22 | 19 | Sand Clay Loam |
| *Ensete ventricosum* | 6.78 | 654 | 2.15 | 0.185 | 10.6 | 51 | 28 | 21 | Sand Clay Loam |

### 3.4 AMF Community abundance

This study which has been conducted in Hawassa area districts of Sidama rigional state uncovered the presence of 1046 spores of 23 different types of AMF morpho-species in 12 AMF genera (Table 3 & 4). The genera are: *Acaulospora, Cetraspora, Claroideoglomus, Dentiscutata, Diversispora, Funneliformis, Gigaspora, Glomus, Racocetra, Rhizophagus, Sclerocystis, and Scutellospora*. Correspondingly, the study conducted in South Ethiopia, Sidama area at agroforestry practicing lands of Shebedino & Wensho districts by [14] had indicated the presence of about 29 morpho-species belonging to 9 genera (*Acaulospora, Glomus, Claroideoglomus*, *Funneliformis*, *Pacispora, Septoglomus, Rhizophagus, Scutellospora* and *Gigaspora*). Here, we could appreciate about 77.78% of similarity in the type of AMF genera abundant in the two study outputs. It might be attributed due to the similarity in physicochemical (moderately acidic and loam soil) feature of the studied areas, regardless of the difference in the type of crops studied.

**Table 3:**
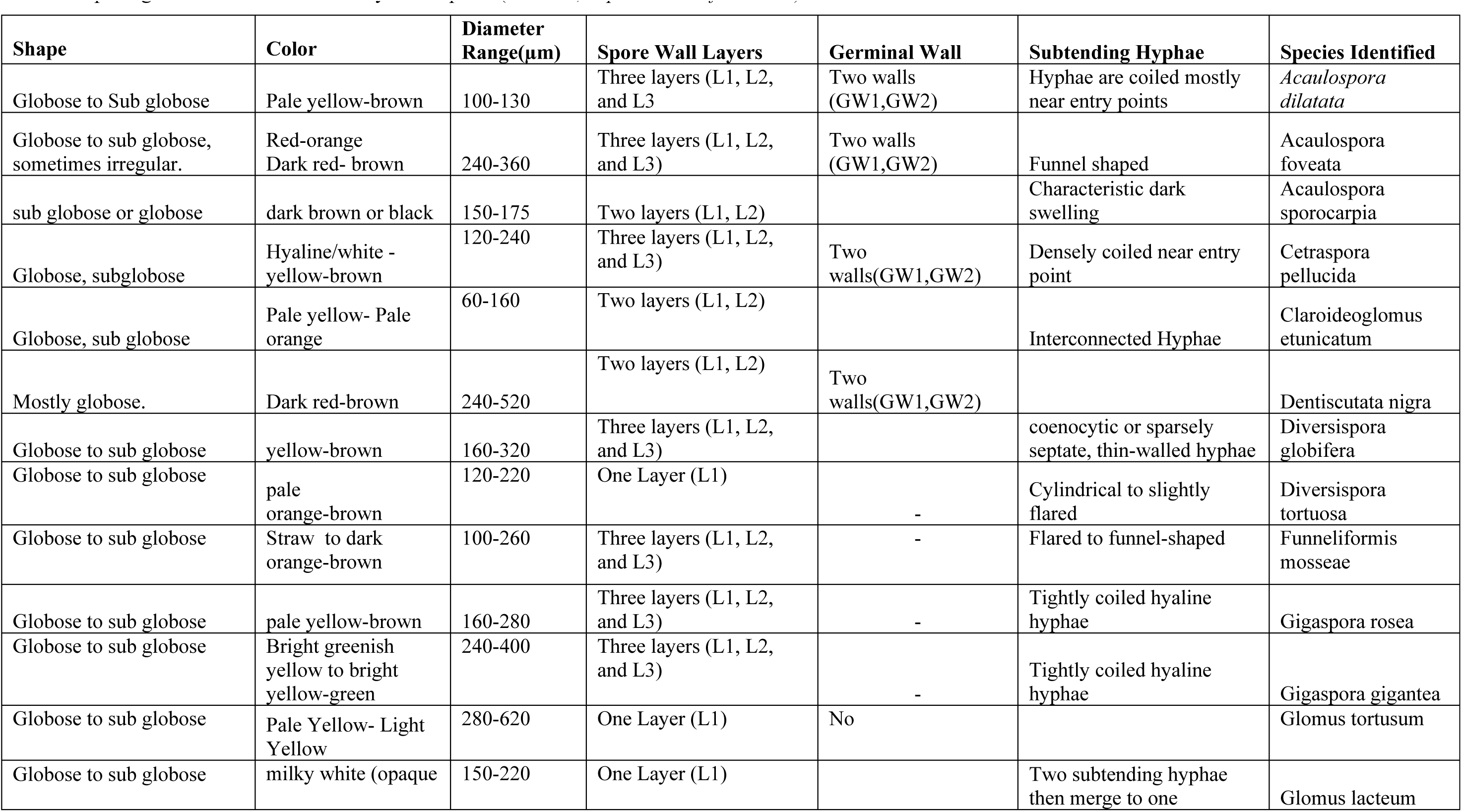
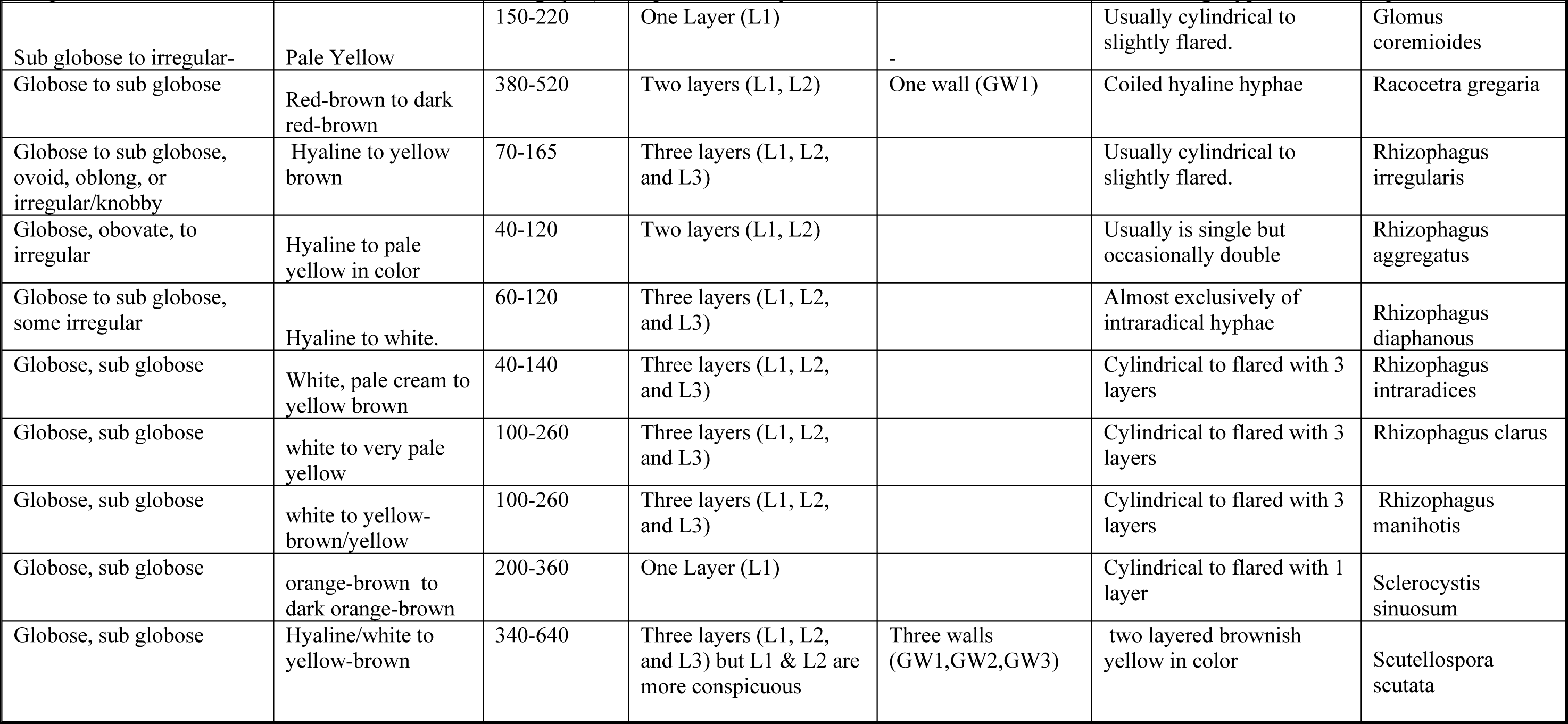
Morphological features used to identify AMF spores (INVAM, http://invam.caf.wvu.edu.)

In another study done in Ethiopia by [24] a total of 42 AMF morpho-species belonging to 15 genera were identified. Similarly, the AMF diversity study in13 selected crops in Sudan [25], in which 42 AMF species belonging to 12 genera were discovered. As we compare the similarity of the current study in the abundance of AMF genera, there is about 83.33% of resemblance with that of [24], it most probably attributed due to the similarity in some important soil physicochemical characteristics (Similarity in soil pH, available phosphorus, and total nitrogen level). However, the land cover of the study by [24] and that of the present one have shown few similarity; nonetheless in both studies land covers such of maize, coffee, tomato,Open Grass Lad were observed.

The genera *Glomus, Funneliformis, Claroideoglomus, Acaulospora, Diversispora, Gigaspora, Rhizophagus, Racocetra, Sclerocystis,* and *Scutellospora*were detected in this study and that of [24]. However, the similarity in AMF genera abundance of this study with the findings in Sudan White Nile State was only in 41.67%. It may be attributed due to the difference in the soil physicochemical feature and the studied crop types. The texture of Hawassa vicinity soil of the specified studied site had not more than 46% clay, whereas that of the White Nile in Sudan had 60% of Clay, pH of the current study soil was neutral, with very low available phosphorus & total nitrogen unlike that of the Sudanese, which was alkaline, very low in available phosphorus, yet highly enriched with total nitrogen. Moreover, out of the 13 studied crops in Sudan, only mango and banana are similar with the present study, whereas the remaining 11 crop lands are not similar with that of the current.Common genera *Glomus, Funneliformis, Claroideoglomus, Acaulospora, and Diversispora* have been discovered instudyoutput of [25] and that of the present one. In general we may conclude that AMF crop association may not always be specific as there is a need to consider other governing parameters such as the physicochemical nature of the soil.

### 3.5 Species richness and Diversity of AMF

Regarding the occurrence of a particular AMF species in certain particular land cover, which is merely termed as spore richness, the monocrop maize has found to be the predominant majorperennial annual crop of all the studied land covers (Table 5). Under the rhizosphere soil of Maize, 8 different types of AMF species were discovered. Following maize permanent crop Enset has taken the second dominance place with 7 different types of AMF species; it again followed by coffee (*Coffea arabica*), Wanza (Cordia africana) and Banana (*Musa acuminat*) with 6 different types of AMF species. The least species richness has been recorded in tomato plant growing in inorganic means, with species richness of only 3 different types of AMF species. Extensive use of agrochemicals might be the cause for such compromised species richness in inorganically cultivated tomato. As this is beyond the scope of this paper, there might be a need to further study the implication of agricultural inputs on the microbial community underneath the soil.

In general, extensive uses of inorganic agricultural inputs have negative impact on AM association. Soils in the conventional agricultural system are AM fungi-impoverished, particularly with regards to number of species [26, 27]. Management practices typical of conventional high input systems, particularly P fertilizer application and the use of biocides, are known to be deleterious to AM fungal symbiosis [28, 29, 30, 31]. This study output corresponds with the findings of [12], that stated AMF species diversity was much lower in tree-based cropping system (Eucalyptus) or mixed croton+juniperus plantation than in the annual cropping systems (monocrops, mixed crops).Similarly, study findings of [24] had reported that, higher species richness of about 31 was obtained in mixed fruit crops ( Avocado, Mango, Coffee, Papaya, Banana, Tomato & Lemon) and 23 mixed Arable land (Teff, sesame, sunflower) compared with the other systems, such as mono-crops with high agricultural inputs of Sorghum, Maize& Natural forest. The results clearly denote that organic management and diversification of crops enhances AMF diversity in low-input agricultural systems.

The diversity of AMF species across the different land use types has also been determined (Table 6). In this regard the diversity of AMF in the rhizosphere soil of the monocrop maize was found to be tremendous of all the crops, with diversity Shannon Weiener Index (H’) of 0.793 ( Table 6) followed by Enset (0.739). However, the diversity of AMF in Eucalyptus, Open Grass Land and Shewshewe has shown similar diversity pattern, with Shannon Weiener Index (H’) of 0.678. The rhizosphere soil under the maize crop has demonstrated a difference of about 80.30% more diverse species of AMF compared to the inorganic input using tomato plantation. The diversity of AMF in inorganically cultivated tomato was recorded to be the lowest of all sorts of land uses; this study has indicated that it was even lower than that of organically growing tomato by about 20.47%. In general infectivity and diversity of AMFCommunities is often reduced in disturbed habitats, where there is extensive use of agrochemicals and various agricultural inputs or post-mining sites [8]. In the current study the lowest diversity index recorded (H’=0.440) was recorded in inorganically cultivated tomato. This finding has been substantiated by other studies. [32] has described that agronomic practices such as monoculture cropping, ploughing, or fertilization have frequently been observed to have a negative impact on the amount as well as the diversity of AM fungi present in soils.

### 3.6 Biomass of AMF in the rhizosphere soil

The biomass of AMF species in the rhizosphere soil across the various land use types has also been determined, through measuring the spore density per 100gm of the sampled soils.In that the AMF under the rhizosphere soil of Eucalyptus tree has shown the highest biomass compared to all other land uses, with spore density of 1907.4±0.404 spores 100g^-1^ of soil (Table 4). The lowest AMF spore density has been recorded in the rhizosphere soil of inorganically cultivated tomato, with spore density of 375.7±0.000 spores 100g^-1^ dry soil. Correspondingly, the study by [33] had described that spore density of the different cropping systems varied significantly within and between land use types ranging from104 spores/100gm soil from Eucalyptus *(E. globulus)* mono (tree) cropping to 929 spores/100gm soil for mixed cropping system (cabbage+sunflower+maize).

**Table 4:** Recovered AMF genera from the rhizosphere soils of different land covers, Hawassa Ethiopia.

| No | Genus | Monocrops |
| --- | --- | --- |
| 1 | Acaulospora | Maize, Enset, Open Grass Land |
| 2 | Cetraspora | Maize |
| 3 | Claroideoglomus | Eucalyptus, Open Grass Land, Wanza,Mango,Pepper |
| 4 | Dentiscutata | Enset,Tomato Organic |
| 5 | Diversispora | Maize,Coffee |
| 6 | Funneliformis | Open Grass Land,Eucalyptus,Wanza,Pepper |
| 7 | Gigaspora | Tomato Inorganic Enset,Tomato Organic,Wanza,Shewshewe |
| 8 | Glomus | Eucalyptus,Wanza,Banana,Mango,Pepper |
| 9 | Racocetra | Coffee |
| 10 | Rhizophagus | Maize,Coffee,Enset,Wanza,Tomato Organic,Banana , Shewshewe, Mango |
| 11 | Sclerocystis | Tomato Inorganic,Maize,Enset,Tomato Organic |
| 12 | Scutellospora | Open Grass Land, Eucalyptus, Wanza, Mango, Pepper |

Study findings in the Southern Ethiopia of Sidama area by [34] has indicated that, the highest number of AM spore population was recorded in rhizosphere soils of *Croton macrostachyus (Bisana)* (1066±19.33) and *Catha edulis (*Khat)(1054±53.12) and the lowest spore density was recorded for *Dioscorea alata (*Boyna)(100.00±2.89) spore per 100 g of dry soil. Omitting the probable similarities and differences in the edaphic factor of the two study areas (The current findings and that of [34] the spore density of the similar land covers can be compared. In that, the output of this study area has shown that; *Ensete ventricosum* (Enset), *Coffea arabica* (Coffee), *Zea mays* (Maize), *Mangifera indica* (Mango) & *Cordia africana* has spore densities of 867, 404.6, 876.63, 549.1, and 1522.07/100gm of soil respectively.

Likewise, that of [34] recorded the spore density in the aforementioned land covers as 630, 995, 700, 580, and 880/100gm of soil respectively; this in general has shown discrepancies between the study outputs, which may be attributed to the physicochemical and management factors of the crops. Unlike the findings of [35], who reported the number of spores produced by AMF in all rhizosphere soils of coffee forests as ranging from 578 to 1313 spores/100g of dry soil the output of this study has indicated that the spore density of coffee to stand at 404.6/100gm of soil. The result of the present study has revealed that mono-cropping and use of biocides do negatively attribute to the reduction in the AMF biomass and limitation of AMF diversity in the soil. More over the physicochemical features of the soils do play a role in the density & diversity of AMF; as the AMF biomass and diversity have demonstrated to differ even with in similar monocrops.

### 3.7 Relative abundance (RA), Isolation frequency (IF) and Importance value (IV)

The relative abundance (RA) literally refers to the spore production potential of AMF. In this regard the current study reported that the top six genera in declining order of spore production from the total recorded 1046 spores as: *Glomus* (18.83%), *Funneliformi s*(13.10%), *Acaulospora* (12.24%), *Claroideoglomus* (12.05%), *Scutellospora* (11.7%), and *Rhizophagus* (10.04%) (Figure 1). The output of the present study was substantiated bythe study findings by [36,37]; in that *Glomus and Funneliformis spp*. had been indicated to have a high spore producing capacity in a shorter time than other genera such as *Gigaspora* and *Scutellospora.* These species could therefore, be selected for future studies as AMFinocula after conducting further testing in their compatibility with different crops and checking their persistence in the field. These AMF genera would be an alternative microbiological technology inputs to high cost requiring and environmentally unfriendly inorganic input crop production, particularly of horticulture production in Ethiopian.

**Figure 1:**
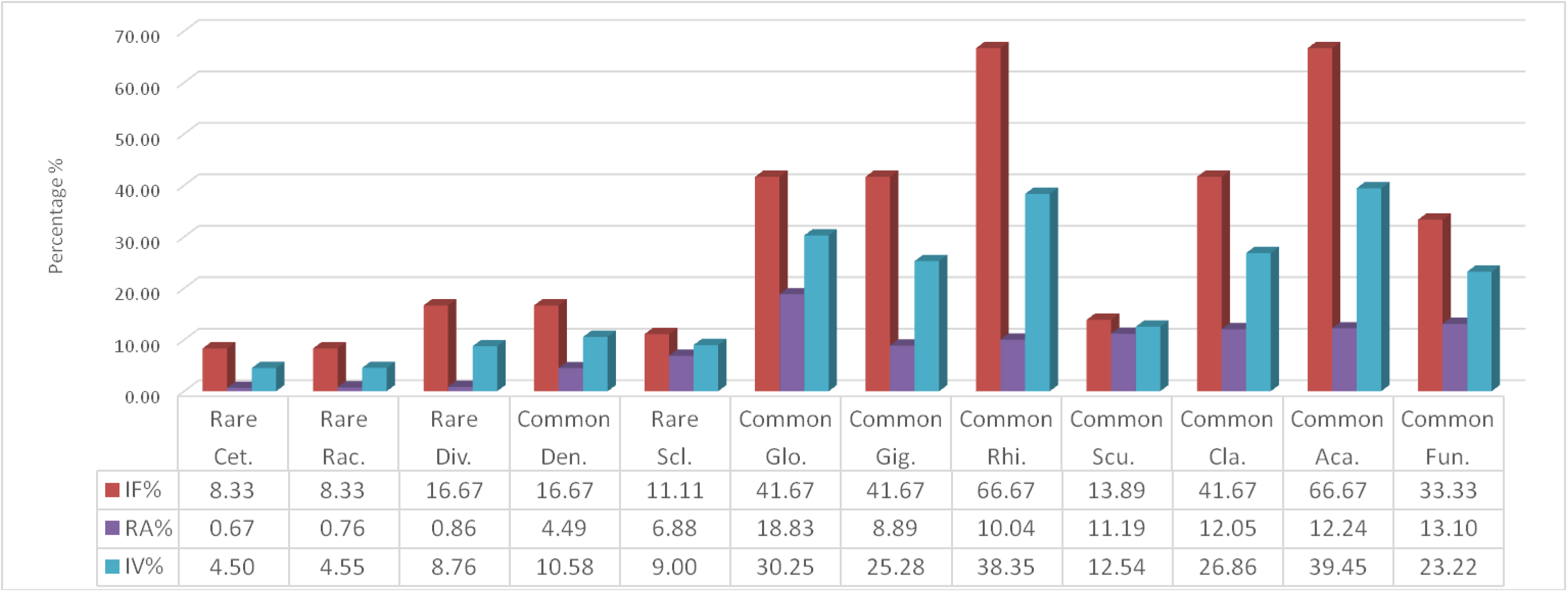
AMF Community Structure (genera level) in the rhizosphere soils of the studied Land Covers, Hawassa Ethiopia. **Key:** Aca. = Acaulospora; Cet. = Cetraspora; Cla. =Claroideoglomus; Den. =Dentiscutata; Div. = Diversispora; Fun. = Funneliformis; Gig. =Gigaspora; Glo. =Glomus; Rac. =Racocetra; Rhi. =Rhizophagus; Sci. =Sclerocystis; Scu. =Scutellospora

Meanwhile, study by [14] showed that spores of four genera *Rhizophugus*, *Glomus, Funneliformis, and Acaulospora* had higher spore production, accounting for 36.22%, 21.20%, 19.39%, 17.54% and 11.74% of the total number of spores respectively. The high spore producing AMF genera lists of [14] has showed the general similarity with the present one; except variation in proportion, as the total spore produced and AMF genera identified are also different. Another study, by [24], in humid low land areas of Ethiopia across the various land uses had indicated the occupancy of larger spore production potential to be concurred by the three genera as; *Claroideoglomus (34.7%) Funneliformis (21.60%) and Glomus (16.80%);* which has a similarity with the current study except the issue of proportional share and order of capacity.

In a nutshellthere is a need toinvestigate these large spore producing AMF genera; *Glomus Funneliformis, Acaulospora*, *Claroideoglomus, Scutellospora*, and *Rhizophagus;* for their AMF inoculum potential; carrying out a through compatibility study against different soil physicochemical feature and crop types. This finding may contribute a bit to shade light for Integrated Pest Management (IPM) endeavors to Ethiopian agricultural system, through using microbiological techniques. As AMF inoculum has multifaceted returns in the strive of crop production; increasing yield and its counteraction against biotic and abiotic factors.

Regarding the isolation frequency of AMF genera, *Acaulospora* and *Rhizophagus* were found to be the most abundantly recovered genera of AMF, which were isolated in 66.67% of the rhizosphere soils investigated. These two dominantly occurring genera were recovered in the rhizosphere soils of *Zea mays*, *Coffea arabica*, *Ensete ventricosum*, *Cordia africana*, *Solanum lycopersicum* (organically growing), *Musa acuminata*, *Casuarina equistifolia*, *Mangifera indica* and Open Grass Land. The second most abundantly isolated genera of AMF were *Claroideoglomus*, *Gigaspora and Glomus*, with isolation frequency (IF) of 41.67%. These were recorded from soils of Eucalyptus, Open Grass Land, Wanza, Mango, Pepper, Tomato (Inorganically & organically growing), Enset, Shewshewe and Banana. Meanwhile, the AMF genera *Funneliformis*has been the third most isolated one, with frequency of encounter in 33.33%of the rhizosphere soils examined. *Funneliformis* were recovered in soils from Open Grass Land, Eucalyptus, Wanza and Pepper.

*Dentiscutata* and *Diversispora* were AMF genera isolated in rhizosphere soils of Enset, Tomato (Organically growing), Coffee and Maize; with isolation frequency of 16.67%. *Racocetra* and *Cetraspora* have been the least frequently isolated AMF genera, which have been encountered in only 8.33% of the soil samples of Maize and Coffee.Similarly, the diversity study of AMF in humid low land areas of Ethiopia by [24] has informed that *Claroideoglomus* (IF=92.9%), *Funneliformis(*IF=91.3%) *and Glomus* (IF=80.2%) had represented the top three AMF genera which encountered frequently from the rhizosphere soils of the various land covers (Teff, Sesame, Maize, Sorghum, Coffee, Acacia & Fruit Crops) studied. Meanwhile, the same study of [24] had reported that*Racocetra* (IF=8.3%), *Entrophospora* (IF=8%) and *Pacispora* (IF=7.7%) as genera with less frequency of encounter. *Claroideoglomus* and *Funneliformis* were dominant genera according to [22], because they were found in all land use types.

The Importance Value (IV) of the genera entails something about the dominancy of the genera. In this regard the current study notified *Acaulospora, Claroideoglomus, Dentiscutata, Funneliformis, Gigaspora, Glomus*, *Rhizophagus* and *Scutellospora* were the commonly encountering genera with <10% IV value <50% (Figure 1). In contrast *Cetraspora, Diversispora Racocetra* and *Sclerocystis* were found to be species of rare occurrence, with IV value <10%. According to [24] *Claroideoglomus (IV=63.8%), Funneliformis (IV=56.4)* stated as dominant genera. However, these two genera were found to be commonly occurring once in the current study. Meanwhile genera which belong to *Glomus (IV=48.5%), Paraglomus (IV=15.7%), Rhizophagus (IV=18.9%), Acaulospora (IV=11.6%), and Gigaspora (IV=208%)* were regarded as commonly occurring once. Similarly to the current study, *Diversispora Racocetra* and*Sclerocystis*were found to be species of rare occurrence. However, [24] had also recorded *Septoglomus*, *Pacispora, Ambispora, Entrophospora, Scutellospora* as rarely occurring AMF genera, unlike the current study which has recorded *Scutellospora* as commonly encountering one, but haven’t encountered the other four rarely occurring AMF in total.

The genera *Glomus*, *Paraglomus, Rhizophagus*, *Acaulospora* and *Gigaspora* were categorized as common. It is interesting to note that more than 50% of the genera were classified as rare. Previous reports have also shown that *Glomus* was dominant in other agro-ecological regions of Ethiopia [12, 38]. The genera *Glomus, Funneliformis,* and *Claroideoglomus* were also reported to be dominant in Cameroon [39] and other sub-Saharan regions, In North Côte d’Ivoire [40], in different land use types of Kenya [41], in the Namibia desert [42], in natural and cultivated savannas of Benin,West Africa [43], in selected crops inthe White Nile State, Central Sudan [25] and in temperate agroecosystems in Europe [44]. The current particular study (Figure 1) is in agreement with AMF composition studies done so far in different parts of Africa, hence it entails for the consideration of the dominant & common AMF genera during AMF inoculum development schemes.

### 3.8 Root colonization by AMF

The total root colonization (RLC) of various land covers by AMF ranges from 11.15-85.41% (Table 5). The root from Open Grass Land (OGL) was colonized by AMF at a rate of 85.41%; it wasthehighest of all types of land cover, followed by Pepper (65.93%). It has got quite strong similarity with the report made by [9], in that the highest AM fungal colonization was found in *Acaciaseyal* (67.3%) from open grazing field (OGF) at Zeway.The least AMF colonization in the present study has been recorded in inorganically cultivated Tomato, Mango & Banana which were 12.94%, 11.15% and 12.3% respectively. Similar study done [14] had indicated that the total percentage mycorrhization (RLC) of the three landuse types was in the range of 55.69% (*Ensete ventricosum,* inmono-cropping) to 90.52% (*Coffea arabica,* in agroforestry).

**Table 5:**
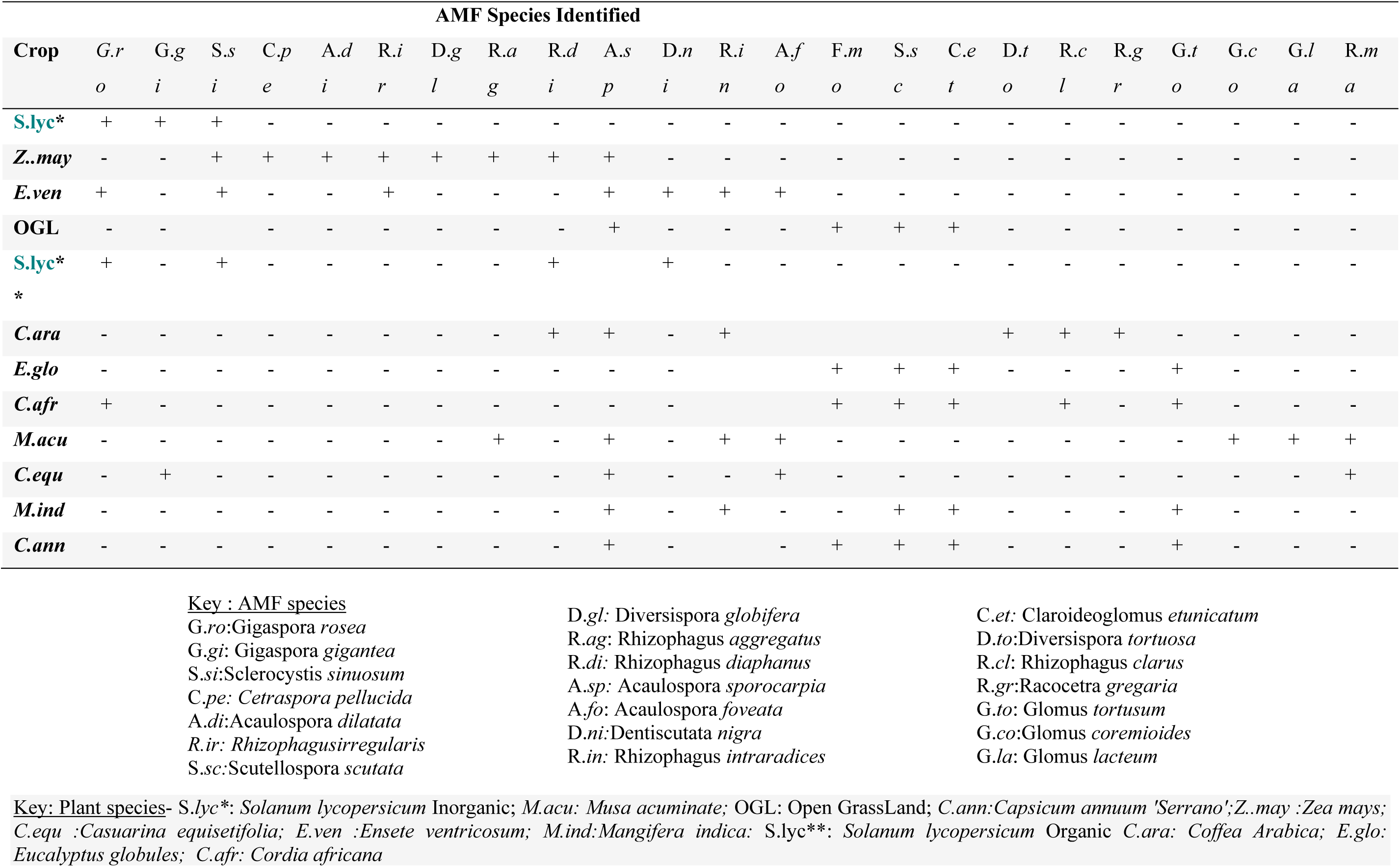
AMF species composition per cropping system, Hawassa, SNNPR.

**Table 6:** AMF species richness and Shannon Weiener index(H’) across different land covers.

| Land Cover | Species Richness | H'(Shannon Wiener Index) |
| --- | --- | --- |
| *Solanum lycopersicum*(Ing) | 3 | 0.44 |
| *Zea mays* | 8 | 0.793 |
| *Ensete ventricosum* | 7 | 0.739 |
| OGL | 4 | 0.530 |
| *Solanum lycopersicum* (Org)) | 4 | 0.530 |
| *Coffea arabica* | 6 | 0.678 |
| *Eucalyptus globulus* | 4 | 0.530 |
| *Cordia Africana* | 6 | 0.678 |
| *Musa acuminata* | 6 | 0.678 |
| *Casuarina equisetifolia* | 4 | 0.530 |
| *Mangifera indica* | 5 | 0.609 |
| *Capsicum annuum* 'Serrano' | 5 | 0.609 |

**Ing-** High Biocide using tomato cultivation; **Org-** No biocides using tomato; **OGL-** Open Grass Land

**Table 8:** Root colonization by AMF across the different Land covers, Hawassa, Ethiopia.

In general, data showed that more mycorrhization occurred in agroforestry (mean mycorrhizal coverage of roots of 71.53%) followed by forest land use systems (with meanmycorrhizal coverage of roots of 68.63%) and the drastically changed mono-cropping system (mean mycorrhizal coverage ofroots of 53.38%). However, with a few exception, annual cropsin mono cropping system showed higher mycorrhization (>80%) than the woody and perennial plants in other land usesystems. This result has shown similarity with that of the current study, in that mycorrhization in monocrops can go beyond (>80%), nonetheless the lowest mycorrhization range have been recorded to go as low as to the level of 11.15% unlike the one reported in South Ethiopia by [14], which is not less than 55.69%.This could be due to the difference in edaphic factors and crop management approaches used between the two study areas. As per the AMF diversity study done in India by Kavitha and Nelson (2013) there had been a discrepancy in mycorrhization rate on the same crop but rationalized as the cause to be edaphic factor. At certain area the total root length colonization of the crop (that was sunflower) had been measured to reach 70% but in the other areas, particularly at marginal soil reach 29% and in fertile soil to reach 35%.In agreement with the aforementioned rationale the findings of root colonization in the present study had a considerable discrepancy from that of the study made by Beyene Dobo*et al.,* 2016b.

To mention the findings in the four similar crops; as per the current study output Wanza, Enset, Coffee & Maize has showed RLC of *14.02; 15.7; 81.1 & 28.53* respectively. Unlike this, the output of [14], had reported that77.93% inWanza, 84.71% in Enset, 90.52 in Coffee & 89.19% in Maize.The AM fungal colonization pattern showed heterogeneity among the roots of the cropping types. As per the study output of [12] the highest hyphal colonization of 73.4% was recorded from sunflower/maize/ (mixed crops) followed by mixed crop (Faba bean+cabbage) 63.4% and pepper (monocrop) with hyphal colonization of 60.3%, correspondingly the present study has reported a root colonization of 65.9 % in Peeper.

The least root colonization was recorded from teff (*Eragrostis tef*) and Eucalyptus monocrops with root colonization of 22%. (P=0.011) and in the current study the lowest root colonization rate which is 11.15% was recorded in Mango, nonetheless the colonization in Eucalyptus was 34.45%.On the contrary, other studies showed that teff and Eucalyptus tree had mycorrhization rate of 58-67% (Cesra *et al.,* 2009) 31-60% [45], respectively. However, lower rate of root infection of Pepper (10-20%) was reported by Castillo *et al.,* (2013). The root colonization of Maize was 73.4% in the study by [12] and 40-44% as per the reportby [46], but in the present study it was recorded as 28.53%, quite different from both reports.

In all the investigated land covers of the present study, the three important structures of the mycorrhizae (arbuscular, vesicular& hyphal-only) were identified. Nonetheless, according to the report by [24] there was no vesicular mycorrhization in low agricultural input using Sorghum. More over as per the study report of [14] there was no Arbuscular and Vesicular colonization in the root length of Wanza, Bisana, Birbira, Korch & Tikurinchet. Meanwhile, as to the report of the current studyorganically cultivated tomato was higher inrate of colonization by either of the AMF structures (Arbuscules, Vesicles & Hypha-only) and total colonization (RLC) compared to the inorganically growing one.

This finding coincided with studies done so far, in that modern agricultural practices such as fertilization, biocide application, and monoculture affect the community composition and diversity of AM fungi [47, 48, 49]. Additionally, although the effect of biocides on AM symbiosis is complex and not easily predictable, overuse of most biocides reduces AM fungi colonization rates and spore production [50, 51].

As per the output of the current study, arbuscular and vesicular colonization of AMF have shown strong and statistically significant positive correlation (r=0.821, p<0.001). Correlation analysis showed that arbuscular colonization was positively correlated with hyphal and vesicular colonization [52]. However the correlation of both arbuscular and vesicular structures to hypha is slight but not statistically significant positive correlation with (r=0.114 P<0.515) and r=0.060, P<0.748) respectively.

There have been spore density and vesicular (r=0.066, P<0.725) as well as arbuscule and spore density (r=0.178, P<0.299) positive correlation, yet it was a slight and not statistically significant one.No significant correlation between AMF colonization and spore density was observed when land-use types were either considered separately or together, which is consistent with several previous reports [14, 53, 54, 55, 56].

## 5. Conclusion

The AMF diversity across the different cropping systems has showed discrepancy. Annual crops demonstrated the largest association with wide varieties of AMF species, followed by permanent crops. The least AMF-crop association was recorded in cropping system where there was use of inorganic agricultural inputs. This signifies the benefit of organic farming to keep the belowground AMF diversity. Acaulospora and Rhizophagus were found to be the most abundantly recovered genera of AMF, which were isolated in 66.67% of the rhizosphere soils investigated. Racocetra and Cetraspora have been the least frequently isolated AMF genera, which have been encountered in only 8.33% of the soil samples. Though not statistically significant, it has been uncovered that there is a positive association between spore density and species richness with that of percentage of OC and total Nitrogen in the rhizosphere.

